# Transfer Entropy Based Causality From Head Motion To Eye Movement

**DOI:** 10.1101/2021.03.11.434910

**Authors:** Zezhong Lv, Qing Xu, Klaus Schoeffmann, Simon Parkinson

**Affiliations:** College of Intelligence and Computing Tianjin University, Tianjin, China; Institute of Information Technology Alpen-Adria Universitat Klagenfurt, Klagenfurt, Austria; School of Computing and Engineering University of Huddersfield, Huddersfield, UK

**Keywords:** coordination of head and eye, unidirectional causality, transfer entropy, behaviometric

## Abstract

Eye movement behavior, which provides the visual information acquisition and processing, plays an important role in performing sensorimotor tasks, such as driving, by human beings in everyday life. In the procedure of performing sensorimotor tasks, eye movement is contributed through a specific coordination of head and eye in gaze changes, with head motions preceding eye movements. Notably we believe that this coordination in essence indicates a kind of causality. In this paper, we investigate transfer entropy to set up a quantity for measuring an unidirectional causality from head motion to eye movement. A normalized version of the proposed measure, demonstrated by virtual reality based psychophysical studies, behaves very well as a proxy of driving performance, suggesting that quantitative exploitation of coordination of head and eye may be an effective behaviometric of sensorimotor activity.

## I. Introduction

Eye movement in natural behavior is important for human environment interaction [1], and provides a fundamental path for visual information acquisition and processing, especially in sensorimotor tasks such as walking and driving [2]. As has been largely accepted, sensorimotor task is executed as a goal-directed activity [3], and the basic mechanism of eye movement during performing sensorimotor tasks results from the coordination of head and eye in gaze changes [4]. Importantly, a specific coordination of head and eye, namely the head-eye coordination is employed in this paper, to indicate the coordination dynamics of head motions preceding eye movements. It is noted that, this specific head-eye coordination mainly contributes to the performing of goal-directed sensorimotor tasks [5].

Usually, eye movement and head motion data are observed as time series of gaze points 〈*X_t_*〉 and of head poses 〈*Y_t_*〉 respectively, labelled by a sequential index *t* =…, 1,2,…. In this paper, stochastic processes are introduced to model the time series of eye movement and of head motion, denoted by *X* and *Y*, respectively, as usually used as natural models for complex and real-world data [6]. In this case, in the procedure of performing sensorimotor task in a dynamic environment, the observation of eye movement at time *t X_t_* naturally depends on the previous *X*_*t*-1_ in probability. That is, the observation of eye movement *X_t_* can be predicted by its past *X*_*t*-1_ in terms of probability [7]. Note that, the head-eye coordination mentioned above indeed points out that, the observation of eye movement *X_t_* is dependent on the past of head motion *Y*_*t*-1_ to some degree. Thus, we consider that, this dependence underlying the head-eye coordination actually indicates that, the predictivity of the current eye movement *X_t_* is added by the past of head motion *Y*_*t*-1_. Notice that, actually, the quantitative causality from one stochastic process *Y* to another *X*, according to its definition, in effect measures the predictivity of *X_t_* added by *Y*_*t*-1_ [8]. As a result, we believe that the head-eye coordination in essence indicates a kind of causality and, we propose a philosophy that this coordination can be quantitatively characterized by a head-eye unidirectional causality, transferring from head motion to eye movement. In this paper, transfer entropy involved in Shannon information theory [6], is utilized as the causality measure, in order to well cope with the complex and non-linear time series of eye movement and of head motion. To go further, the visual cognitive processing state, which corresponds to the head-eye coordination [5], has a direct correlation with the performance of goal-directed sensorimotor task [9]. In this case, we suppose that the strength of the specific head-eye coordination, which is obtained based on the unidirectional causality from head motion to eye movement, shoud indicate the task performance.

Notice that, although the research on the patterns of the coordination of head and eye has attracted a lot of attention recently [4] [5], there is no quantitative investigation on this coordination for the sake of deep theoretical understandings and of behaviometric applications. To fill this gap, we success in obtaining two new quantitative causality measures for representing this coordination and driving performance, respectively. It is worthy to indicate that the proposed transfer entropy based causality measures provide a new window into the discovery of behaviometric. The main contributions of this paper are listed as follows:

- A transfer entropy based head-eye unidirectional causality measure (*UCM*) is proposed to quantitatively evaluate the specific coordination of head and eye, this coordination involves head motions preceding eye movements.
- A normalized unidirectional causality measure (*NUCM*) is proposed for the purpose of being quantitatively compatible with task performance. This *NUCM* measure works well as a proxy of driving performance, verified by our psychophysical studies conducted in this paper.

## II. Related Works

For the sake of this paper, the concepts of Wiener causality [8] and of transfer entropy [6] are reviewed.

Wiener causality is one of the most common causality measures [8]. The abstract idea of Wiener causality is that, for two random variables *P* and *Q*, if the past of *Q* adds the predictivity of current *P*, then it is said that there is a causality from *Q* to *P*.

Transfer entropy is a well-known way to measure the causality between time series [6]. Transfer entropy is more general in the sense that it is capable of handling complex and non-linear time series, compared with classic causality measures. Given random variables *P* and *Q*, transfer entropy from the source *Q* to the target *P* is defined as follows [6]

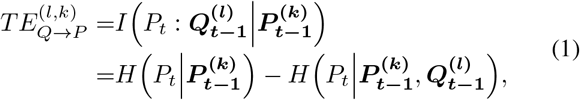

where *P_t_* and *Q_t_* are the observations of variables *P* and *Q* at time *t* respectively, 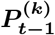 and 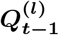 are the histories of target and source variables {*P*_*t*-1_,…,*P_t-k_*} and {*Q*_*t*-1_,…, *Q_t-l_*} respectively, and *H*(·|·) means the conditional entropy. Here *l* and *k* are the so-called history lengths of *Q_t_* and of *P_t_* respectively.

Obviously, transfer entropy is asymmetric. Because 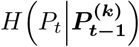 is no smaller than 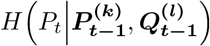, transfer entropy is nonnegative. And considering that 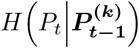 and 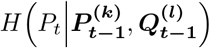 are nonnegative, the maximum of 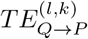 takes 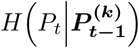.

## III. Experiment

### A. Task

Driving, which is commonly executed as a goal-directed activity, is taken as the sensorimotor task in our psychophysical experiments. In this paper, the driving task consists of maintaining the driving speed at 40 km/h. In this paper, the inverse of the mean acceleration during driving is taken as the indicator of driving performance, as usually used in literature [10]. That is, the larger the mean acceleration is, the worse driving performance becomes, and *vice versa*.

### B. Apparatus

The psychophysical experiments in this paper are conducted in a virtual reality environment through the display by a *HTC Vive* [11] headset. And there is a *7INVENSUN Instrument aGlass DKII* eye-tracking equipment [12] embedded in the headset. An illustration of the headset with the embedded eye tracker is given in Fig. 1. Eye movement (gaze point) and head motion (head pose) data are recorded in a frequency of 90 Hz by the eye-tracking equipment and by the headset respectively. Head motion and eye movement are both captured as pitch and yaw [5]. Virtual driving is performed based on a *Logitech G29* steering wheel [13]. A desktop monitor is utilized to observe the eye movement and driving behaviors of participants in the procedure of experiment.

**Fig. 1.**
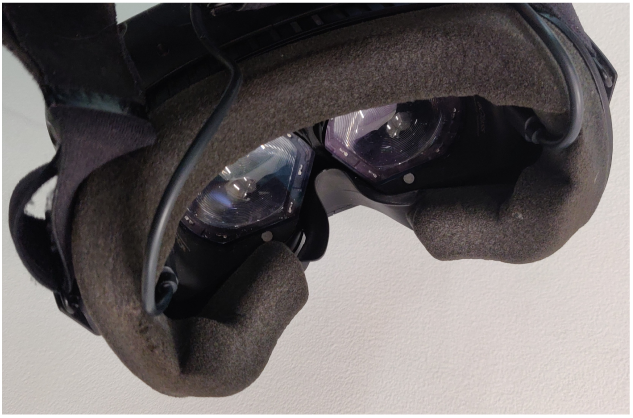
A *HTC Vive* headset and a *7INVENSUN Instrument aGlass DKII* eye tracker.

### C. Virtual reality environment

Virtual reality provides an effective way to investigate eye movement and driving [14]. In this paper, the virtual environment for the psychophysical studies consists of common roads and buildings, and without visual stimuli based distractors such as traffic lights and pedestrians. This is because that, the specific head-eye coordination is mainly contributed by topdown modulation. Example illustrations of performing driving task and of the virtual environment are presented in Fig. 2.

**Fig. 2.**
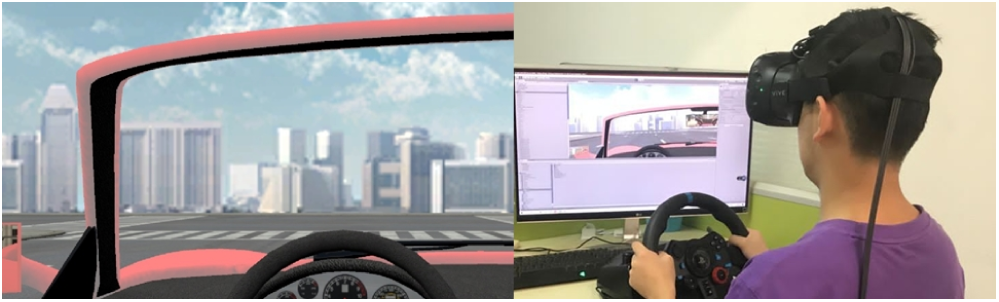
A participant is performing the driving task in the virtual environment.

### D. Participants

Twelve participants (7 male, 5 female; ages 22.9 ± 1.95), with normal color vision and normal/corrected-to-normal visual acuity, participate in the psychophysical study. All of the participants hold their driver licenses for no less than one and a half years. And none of the participants has adverse reaction to the virtual environment utilized in this study.

### E. Procedure

The purpose and procedure about the psychological studies are introduced to the participants through a preparation session.

Four test sessions, which involve the same driving task and the same driving routes, are completed by each participant. For each participant, there is an interval of three days between every two test sessions. Before every test session, a 9-point calibration for the eye tracker is applied for the eye tracker. In this paper, we use a trial to denote a test session. And there are 12 * 4 = 48 valid trials obtained by the psychophysical experiments.

## IV. Methodology

### A. An unidirectional causality measure

As has been discussed in Section I, the head-eye coordination can be described as an unidirectional causality measure (*UCM*) from head motion to eye movement and, transfer entropy is selected to investigate this causality due its generality. Following (1), transfer entropy from head motion *Y* to eye movement *X*, *TE_Y→X_*, is defined as

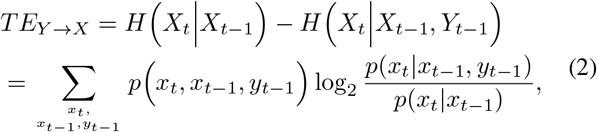

note in this paper, the history lengths of *X* and *Y* are both taken as one, as usually done in literature [6]. Here *p*(·), *p*(·|·) denote the (conditional) probabilities of distributions of gaze and head data. And similarly, *TE_X→Y_*, transfer entropy from eye movement to head motion, is given as follows

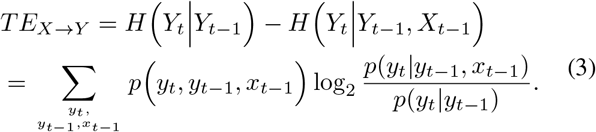

Notice that, the larger the causality from *Y* to *X*, the more predictivity of current *X* added by the past of *Y*, the larger *TE_Y→X_* is; and analogously this also applies to *TE_X→Y_*. In this case, the unidirectional causality from *Y* to *X* can be effectively investigated by the difference between *TE_Y→X_* and *TE_X→Y_*,

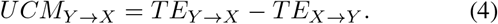

Importantly, the proposed *UCM_Y→X_* establishes a definition of cause-effect relationship between head motion and eye movement. *UCM_Y→X_* > 0 represents that head motion is the cause and eye movement is the effect, *UCM_Y→X_* < 0 indicates the causality from eye movement to head motion, and *UCM_Y→X_* = 0 (practically *UCM_Y→X_* is small enough) means there is no unidirectional causality.

A hypothesis test is performed for determining whether there exists a valid *UCM_Y→X_* with a high confidence level. To do this, the null hypothesis *H*_0_ is that *UCM_Y→X_* is small, that is, it means that *X* and *Y* do not influence each other. To verify or reject *H*_0_, surrogate time series 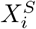 and 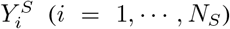 of the original *X* and *Y* are used. For surrogate generation, random shuffle is done because the history lengths of *X* and *Y* are both taken as one [6]. The unidirectional causality from the 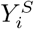 to 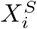, following (4), is obtained as follows

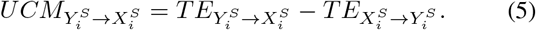

Then the significance level of *UCM_Y→X_* is defined as

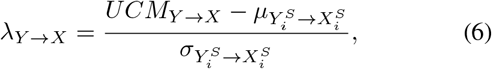

where 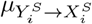 and 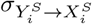 are the mean and standard deviation of 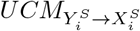, respectively. The probability to reject *H*_0_ can be obtained based on Chebyshev’s inequality

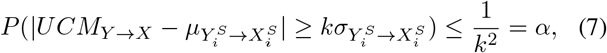

where 1 – *α* is the confidence level to reject *H*_0_, and parameter *k* is any positive real number. The number of the surrogates, which is related with the confidence level, is obtained as

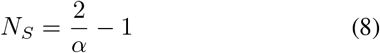

for a two-sided test.

Usually, that the parameter *k* used in (7) is taken as 6 results in a confidence level of 97.3% [15]. This means that, if the significance level is bigger than 6 (*λ_Y→X_* > 6), then, equivalently with a confidence level of more than 97.3%, there exists a head-eye unidirectional causality from head motion to eye movement. Note that the significance and confidence levels are the same in use in hypothesis test. Additionally, according to statistical test theory [15], it is important to know that a minimum confidence level, acceptable in practice, is 95% (here the corresponding significance level is 4.47).

### B. A normalized unidirectional causality measure

In a goal-directed driving, the head-eye coordination corresponds to the state of visual-cognitive processing [5], and meanwhile this state represents the performance of sensorimotor tasks [9]. As discussed in Section IV-A, the unidirectional causality from head motion to eye movement in effect gives an estimation of the head-eye coordination. Therefore, we propose a hypothesis that the head-eye unidirectional causality should work well as a proxy of the driving performance. Considering that correlation analysis is a classic and popular tool for investigating a linear relationship between variables, this tool is utilized to verify this hypothesis in our paper. Because correlation analysis is actually done through the comparison between variable values [**?**], we propose a normalized unidirectional causality measure from head motion to eye movement, *NUCM_Y→X_*, for being quantitatively compatible with the driving performance, as follows

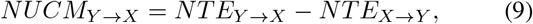

where

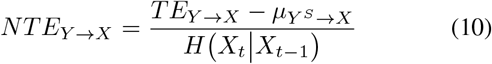

is a kind of normalized transfer entropy [16] used for *TE_Y→X_*, and note this normalization is effective and popularly used [16]. Here *μ_Y^s^→X_* is the mean of the transfer entropies 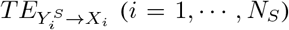 from the surrogate head motion to eye movement, and *H*(*X_t_*|*X*_*t*-1_) is the maximum of *TE_Y→X_*. *NTE_X→Y_* can be obtained similarly.

*NUCM_Y→X_* values from −1 to 1. Here, *NUCM_Y→X_* is, as a matter of fact, a normalized measure of the strength of the specific head-eye coordination. This strength actually reflects the state of visual-cognitive processing in a quantitative way. Thus, in the procedure of performing sensorimotor task, the bigger this strength is, the better the state of visual-cognitive processing becomes, the higher the task performance is; and *vice versa*. It is emphasized that the normalized unidirectional causality measure can be easily and very well exploited in real applications such as safety driving, for providing an effective technique to monitor and obtain the cognitive processing states of drivers. And this exploitation offers a valuable and potential behaviometric for sensorimotor tasks in application scenarios.

## V. Results and Discussions

### A. Transfer entropies between head motion and eye movement

A significant difference between *TE_Y→X_* and *TE_X→Y_*, is revealed by one way analysis of variance (*ANOVA*) (*F*(1,94) = 80.25,*p* < 0.05), as illustrated in Fig. 3. Obviously, *TE_Y→X_* is much bigger than *TE_X→Y_*, as demonstrated in Fig. 3. These computation results clearly verify the specific head-eye coordination. Notice that, our quantitative finding here is accordance with the observation that, during performing sensorimotor tasks, eye movement is done through the coordination of head and eye involving head motions preceding eye movements [5].

**Fig. 3.**
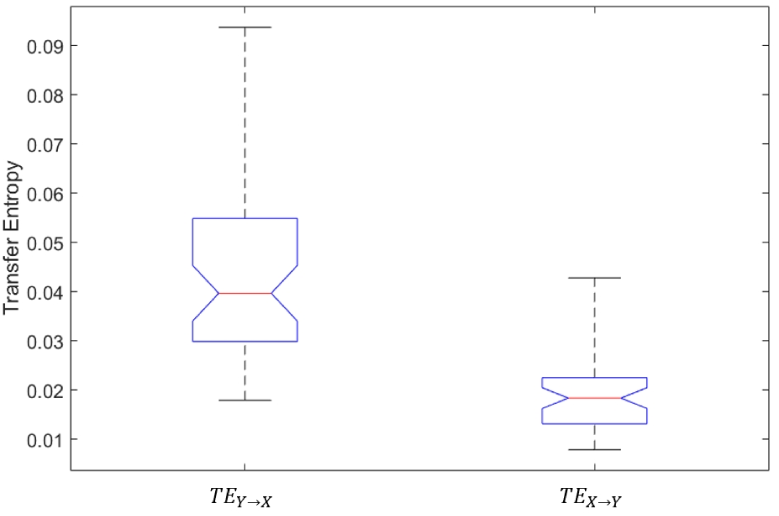
Transfer entropies between head and gaze data.

### B. The unidirectional causality from head motion to eye movement

Based on the transfer entropies between head motion and eye movement, *UCM_Y→X_* is proposed for a deeper understanding of the specific head-eye coordination. Table I gives the *UCM_Y→X_* results obtained in trials by all the participants. The strictly positive *UCM_Y→X_* values discover that there exists, with high confidence (its corresponding results are presented in Table II), an unidirectional causality from head motion to eye movement, in the procedure of performing goal-directed sensorimotor tasks. It is valuable to emphasize that, our creativity, considering the specific head-eye coordination as an unidirectional causality, is strongly verified by this discovery.

**TABLE I.** The unidirectional causality measures *UCM_y→x_* (*10^-2^)

| Participant | 1st Trial | 2nd Trial | 3rd Trial | 4th Trial |
| --- | --- | --- | --- | --- |
| 1 | 0.79 | 0.78 | 0.17 | 1.24 |
| 2 | 1.21 | 4.33 | 1.88 | 2.76 |
| 3 | 1.43 | 0.61 | 0.30 | 1.82 |
| 4 | 1.18 | 0.98 | 1.85 | 2.70 |
| 5 | 3.14 | 4.73 | 6.74 | 2.25 |
| 6 | 3.55 | 4.43 | 3.02 | 1.97 |
| 7 | 2.91 | 3.42 | 4.33 | 3.65 |
| 8 | 0.91 | 2.02 | 3.86 | 1.57 |
| 9 | 3.33 | 4.33 | 3.75 | 1.62 |
| 10 | 0.68 | 3.06 | 1.91 | 3.09 |
| 11 | 2.76 | 1.21 | 1.89 | 1.70 |
| 12 | 2.26 | 5.53 | 6.18 | 7.01 |

**TABLE II.** Significance levels *λ_Y→X_*

| Participant | 1st Trial | 2nd Trial | 3rd Trial | 4th Trial |
| --- | --- | --- | --- | --- |
| 1 | 8.9 | 9.2 | 15.2 | 10.3 |
| 2 | 17.3 | 18.1 | 7.8 | 5.7 |
| 3 | 17.0 | 5.5 | 11.2 | 11.6 |
| 4 | 11.9 | 15.0 | 12.2 | 16.7 |
| 5 | 16.0 | 12.1 | 12.5 | 8.0 |
| 6 | 15.8 | 20.2 | 21.1 | 17.1 |
| 7 | 17.3 | 19.3 | 26.3 | 22.9 |
| 8 | 7.9 | 13.9 | 13.6 | 10.7 |
| 9 | 16.4 | 17.8 | 17.4 | 13.9 |
| 10 | 14.3 | 16.9 | 13.1 | 14.5 |
| 11 | 19.3 | 12.7 | 18.9 | 19.4 |
| 12 | 14.7 | 12.5 | 16.7 | 15.0 |

The significance level *λ_Y→X^s^_* in hypothesis test are presented in Table II. Almost all the values of *λ_Y→X^s^_* are higher than 6, that is, the corresponding confidence levels are more than 97.3%. There are only two exceptional *λ_Y→X^s^_*, 5.7 and 5.5, marked with boxes, are slightly lower than 6. Even here, the corresponding confidence levels are 96.9% and 96.7% respectively and, this is acceptable in statistics for practical use [17].

It is also worthwhile noting that, the head-eye unidirectional causality *UCM_Y→X_* is obtained by hypothesis test based on the use of surrogate time series [15]. Because the history length used in the time series data of this paper is taken as 1 [6], random shuffle is done for obtaining the surrogate time series. As for other possible options of history length, iterative amplitude adjusted Fourier transform (*iAAFT*) [18] would be considered and, this is for future work. Considering that hypothesis test is a necessary computational component to estimate causality measures, hypothesis test by *e-values* [19] would be as future consideration for a more effective solution.

### C. Correlation analyses

A significant correlation (*p* < 0.05) between *NUCM_Y→X_* and driving performance is revealed by the three correlation analyses [20], with Pearson linear correlation coefficient (*PLCC*), Spearman rank order correlation coefficient (*SROCC*) and Kendall rank order correlation coefficient (*KROCC*) of 0.32, 0.41 and 0.27 respectively, as shown in Table III. Additionally, as shown in Fig. 4, this correlation analysis is confirmed by a linear regression, demonstrating a strong linear relationship between *NUCM_Y→X_* and driving performance (*y* = 0.445 + 1.51*x*). Obviously, these results point out that, in the procedure of performing goal-directed sensorimotor tasks, the proposed normalized unidirectional causality from head motion to eye movement, *NUCM_Y→X_*, offers a good indicator of task performance.

**Fig. 4.**
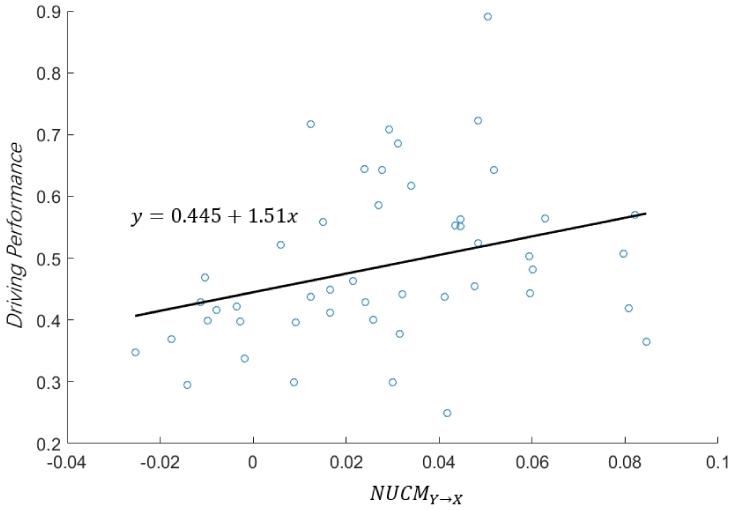
Linear regression analysis between *NUCM_Y→X_* and driving performance.

**TABLE III.**
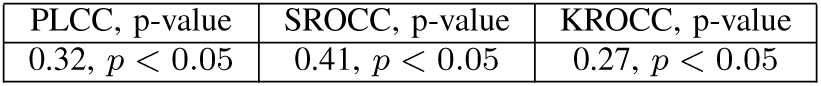
Correlation analysis results

| PLCC, p-value | SROCC, p-value | KROCC, p-value |
| --- | --- | --- |
| 0.32, $p < 0.05$ | 0.41, $p < 0.05$ | 0.27, $p < 0.05$ |

In this paper, a mean acceleration is used as the driving performance [10], and this is in accordance with the task goal, namely maintaining driving speed in our virtual driving based psychophysical studies. It is obvious that the fulfillment of this kind of driving essentially constitutes a goal-directed sensorimotor activity. Notably, it has been observed that eye movement behavior involved in this kind of goal-directed activity is provided through a head-eye coordination in gaze changes, with head motions prior to eye movements [5]. Considering that the proposed transfer entropy based causality behaves as a good indicator of driving performance, and that our causality measure achieves an estimation of the strength for the head-eye coordination, the correspondence between the strength of head-eye coordination and driving performance is thus clearly presented and revealed. Actually, our discovered correspondence is in accordance with the combination of two previous findings. One finding is that there is a direct link between the state of visual-cognitive processing and task performance [9]; the other is that the head-eye coordination represents a state of visual-cognitive processing [5].

Employing normalization on the proposed *UCM* is a necessary way for obtaining a quantitative strength of the head-eye coordination to be compatible with the driving performance [16]. Other normalization methods will be done in future work [21].

In the procedure of performing a goal-directed sensorimotor task, coordination of head and eye is mainly contributed by the top-down modulation, as has been largely accepted in literature [3]. In order to ensure that top-down modulation plays the most important role in the goal-directed driving, we take speed control as the single driving task. And meanwhile, the virtual environment is specially designed to include no visual stimuli based distractors (such as traffic lights and pedestrians). As a result, the proposed *UCM* and *NUCM* can be evaluated and investigated clearly and easily in this paper.

Last but also important, many practical applications can take advantage of our proposed causality measures for the discoveries and applications of behaviometric, as has been emphasized in Section IV-B.

## VI. Conclusion and Future Works

A major progress has been achieved for deeply understanding the head-eye coordination in gaze changes, with head motions preceding eye movements. Two new quantitative measures, namely the unidirectional causality measure and its normalized version, have been presented for estimating the quantification of the head-eye coordination. Based on psychophysical studies conducted, we have clearly discovered that, during virtual driving, the two proposed measures behave very well as indicators for cause-effect link between head and gaze data, and for driving performance, respectively. Our discovery also offers a large potential for behaviometric applications.

In the near future, the unidirectional coordination from eye movements to head motions, which results from stimulus-driven shifts of attention [5], will be investigated for its quantification, by the exploitation of transfer entropy based causality. As has been discussed in Sections V-B and V-C, some more effective approaches for hypothesis test and for the normalization of transfer entropy will be studied.

## References

[1] Mary Hayhoe and Dana Ballard, “Eye movements in natural behavior,” Trends in Cognitive ences, vol. 9, no. 4, pp. 188–194, 2005.

[2] Kosobudzki, Marcin, Jozwiak, Zbigniew, Kapitaniak, Bronislaw, Walczak, Marta, Bortkiewicz, and Alicja, “Application of eye-tracking in drivers testing: A review of research,” International Journal of Occupational Medicine and Environmental Health, 2015.

[3] Kurz Johannes, Hegele Mathias, and Munzert Jorn, “Gaze behavior in a natural environment with a task-relevant distractor: How the presence of a goalkeeper distracts the penalty taker,” Frontiers in Psychology, vol. 9, 2018.

[4] Edward G. Freedman, “Coordination of the eyes and head during visual orienting,” Experimental Brain Research, vol. 190, no. 4, pp. 369, 2008.

[5] A. Doshi and M. M. Trivedi, “Head and eye gaze dynamics during visual attention shifts in complex environments,” Journal of Vision, vol. 12, no. 2, pp. 189–190, 2012.

[6] T Bossomaier, L Barnett, and J T Lizier, An introduction to transfer entropy: information flow in complex systems, Springer Publishing Company, Incorporated, 2016.

[7] Robin S. Weiss, Roger Remington, and Stephen R. Ellis, “Sampling distributions of the entropy in visual scanning,” Behavior Research Methods,Instruments, Computers, 1989.

[8] N. Wiener, The Theory of Prediction, Modern Mathematics for Engineers, McGraw-Hill, 1956.

[9] John H. L. Hansen, Carlos Busso, Yang Zheng, and Amardeep Sathyanarayana, “Driver modeling for detection and assessment of driver distraction: Examples from the utdrive test bed,” IEEE Signal Processing Magazine, vol. 34, no. 4, pp. 130–142, 2017.

[10] Ankit Kumar Yadav and Nagendra R Velaga, “Effect of alcohol use on accelerating and braking behaviors of drivers,” Traffic Injury Prevention, vol. 20, no. 4, pp. 353–358, 2019.

[11] “Htc corporation,” https://www.htcvive.com.

[12] “7invensun instrument aglass,” https://www.7invensun.com/xnxsxqy.

[13] “Logitech g29,” https://www.logitechg.com/en-us/products/driving/driving-force-racing-wheel.html.

[14] Ludwig Sidenmark and Hans Gellersen, “Eye, head and torso coordination during gaze shifts in virtual reality,” ACM Transactions on Computer-Human Interaction, vol. 27, no. 1, pp. 1–40, 2019.

[15] Thomas Schreiber and Andreas Schmitz, “Surrogate time series,” Physica D Nonlinear Phenomena, vol. 142, no. 3, pp. 346–382, 1999.

[16] R. Marschinski and H. Kantz, “Analysing the information flow between financial time series,” Physics of Condensed Matter, vol. 30, no. 2, pp. 275–281, 2002.

[17] James Theiler, Stephen Eubank, Andr Longtin, Bryan Galdrikian, and J. Doyne Farmer, “Testing for nonlinearity in time series: The method of surrogate data,” Physica D-nonlinear Phenomena, vol. 58, no. 1-4, pp. 77–94, 1992.

[18] Sergio Albeverio, Nonlinear Time Series Analysis, Cambridge University press, 1997.

[19] Vladimir Vovk and Ruodu Wang, “E-values: Calibration, combination, and applications,” Annals of Statistics, forthcoming, 2020.

[20] “Correlation and dependence,” https://en.wikipedia.org/wiki/Correlation and dependence.

[21] Xuegeng Mao and Pengjian Shang, “Transfer entropy between multi-variate time series,” Communications in Nonlinear ence and Numerical Simulation, vol. 47, pp. 338–347, 2016.

